# Characterization of the Gut Microbiota and Serum Metabolomics in Patients with Type 2 Diabetes Mellitus and Newly Diagnosed Acute Coronary Syndrome

**DOI:** 10.64898/2026.02.09.704968

**Authors:** Hongliang Wu, Yihui Teng, Ruiyi Chen, Haimi Zhao, Wangang Guo, Kai Wang, Haowei Xu, Jiaying Zhou, Yan Li, Yuerong Xu, Mingming Zhang

## Abstract

Biomarkers for the early identification of acute coronary syndrome (ACS) risk remain inadequately investigated, particularly in patients with type 2 diabetes mellitus (T2DM), for whom timely clinical intervention may substantially enhance prognostic outcomes. The gut microbiota and serum metabolites may serve as pivotal mediators in the occurrence and progression of ACS among patients with T2DM, and whether these factors can be used for the precise discrimination of patients with T2DM complicated by ACS remains to be explored. Overall, 76 participants were enrolled (38 patients diagnosed with T2DM complicated by ACS and another 38 with T2DM without ACS). 16S rRNA sequencing combined with untargeted LC-MS metabolomics revealed a dysregulated gut-serum axis in patients with T2DM complicated by ACS: enrichment of proinflammatory microorganisms (*Enterococcus* spp.), reduction of butyrate producers (*Butyricimonas* spp.) and concomitant dysregulation of circulating lipid metabolites-upregulation of PC(16:0/9:0 (CHO)) and arachidonic acid alongside downregulation of cholesterol sulfate. By integrating multiomics data and applying various feature selection methods, we subsequently identified six key biomarkers. The final constructed combined model robustly distinguished patients with T2DM complicated by ACS from those with T2DM alone (AUC = 0.983), outperforming the other single omics models. Our study revealed that the gut microbiota and related serum metabolites serve as key mediators in the onset and progression of ACS among patients with T2DM, and demonstrated their potential value as noninvasive biomarkers for the early diagnosis of T2DM complicated by ACS.

**Clinical Perspective:** *What Is New?:* We combined 16S rRNA sequencing with untargeted LC-MS metabolomics to dissect the gut-serum axis in patients with T2DM complicated by newly diagnosed ACS. We identified distinct gut microbial and serum metabolic signatures that distinguish ACS progression within the T2DM population. A multiomics classifier integrating clinical, microbial, and metabolic variables achieved robust diagnostic performance (AUC = 0.983) and outperformed single omics models.

*What Are the Clinical Implications?:* Integrated multiomics biomarkers facilitate the early identification of ACS progression in patients with T2DM, offering novel avenues for precision prevention, dynamic clinical surveillance, and individualized therapeutic strategies. Dysregulations of the gut microbiota and serum metabolome may play an important role in the development and progression of ACS in patients with T2DM.

## Introduction

Type 2 diabetes mellitus (T2DM) and acute coronary syndrome (ACS) are two major diseases that currently pose severe threat to human health. According to the statistics from the International Diabetes Federation (2025), approximately 589 million (11.1%) adults aged 20–79 years worldwide have diabetes, showing a continuously significant upward trend, and an additional 252 million patients remain undiagnosed ^1^. Diabetes mellitus acts as a key risk factor for coronary heart disease; approximately 30% of patients with T2DM develop comorbid coronary artery disease (CAD), with ACS being the most common. Compared with nondiabetic patients, the mortality rate of these patients is significantly greater, which significantly decreases their life expectancy and imposes a heavy medical and economic burden on society ^1–4^. ACS is the most severe clinical phenotype of CAD. Its main mechanism involves thrombosis induced by unstable plaque rupture, leading to complete or incomplete occlusion of coronary arteries and subsequent myocardial ischemia, which is a major cause of patient death ^5^. Although the formation and progression of atherosclerotic plaques are usually relatively slow, dysregulations of glucose and lipid metabolism and insulin resistance among T2DM patients can accelerate this process through mechanisms such as oxidative stress and chronic inflammation, resulting in the rapid deterioration of clinical conditions within a short period ^6–8^. In addition, the arterial lumens of patients with T2DM complicated by ACS are usually narrower and often present with multivessel lesions and diffuse lesions. These patients typically have atypical or asymptomatic myocardial ischemia ^9^ and require multiple percutaneous coronary interventions, with a high rate of readmission, a high incidence of heart failure, and poor prognosis ^10,11^.

However, reliable indicators for assessing the progression of T2DM complicated by ACS are lacking in clinical practice. Current diagnostic approaches, including coronary CT angiography and coronary angiography, fail to accurately identify plaque characteristics and thrombus burden. Although high-resolution intracoronary optical coherence tomography and intravascular ultrasound can more effectively distinguish plaque rupture, erosion, coronary nonobstructive myocardial infarction and spontaneous coronary artery dissection, their invasiveness and high cost have limited their ability to detect vulnerable plaques in asymptomatic patients ^12^. This poses a significant challenge to the early identification of ACS in high-risk populations. Therefore, identifying more ideal biomarkers for the early recognition and diagnosis of ACS risk in patients with T2DM to provide better strategies for precision medicine is an urgent need.

The gut microbiota is not only a crucial component of human symbiosis but also an endocrine organ that can produce various bioactive compounds that enter the circulatory system to regulate host health ^13^. In patients with T2DM, altered gut microbiota and serum metabolomics can affect the occurrence and progression of the disease through intestinal barrier impairment and inflammatory responses, with some metabolites being associated with cardiovascular outcomes in patients with T2DM ^14,15^. In cardiovascular diseases, the gut microbiota and their associated metabolites exert critical regulatory effects via the “gut–heart axis” ^16,17^. Changes in serum metabolites and the gut microbiota occur between healthy individuals and patients with CAD at different stages and are correlated with disease severity; combinations of specific bacteria and their mediated metabolites can significantly distinguish patients with different CAD subtypes ^18^. The most critical subtype of CAD, ACS, has been confirmed to possess unique serum metabolomic and gut microbial characteristics, manifested mainly as increased levels of proinflammatory bacteria and circulating metabolites, which decrease after clinical recovery to chronic coronary syndrome. Moreover, a combined model of microbial taxa, metabolites, and clinical biomarkers can accurately distinguish patients with ACS from healthy controls ^19^. For high-risk populations such as patients with T2DM, whether the gut microbiota and its metabolites undergo further alterations following secondary ACS, and whether clinical indicators, the gut microbiota, and serum metabolomics can accurately differentiate these two groups of patients (those with T2DM with ACS vs. those with T2DM alone), remain to be investigated.

We profiled the gut microbiota and serum metabolites in patients with T2DM alone and T2DM complicated with newly diagnosed ACS, and delineated core biomarkers linked to ACS progression. On the basis of multiomics data analysis, we established a classifier that can accurately distinguish patients with T2DM complicated with newly diagnosed ACS from those with T2DM alone. These findings decode gut–metabolite drivers of coronary atherothrombosis in patients with T2DM, enabling precision prevention, clinical surveillance, and personalized treatment.

## Materials and Methods

### Study Design and Ethical Approval

This case–control study consecutively recruited 76 patients at Tangdu Hospital, Fourth Military Medical University. The T2DM-ACS group (n = 38) comprised patients with T2DM complicated by ACS, while the T2DM group (n = 38) comprised patients with T2DM alone.

Patients with confirmed T2DM alone and those with T2DM complicated by newly diagnosed ACS were enrolled in the present study. Detailed inclusion and exclusion criteria are described in the Supplemental Appendix.

This research received approval from the Ethics Committee of Tangdu Hospital, Air Force Medical University (Approval No.: K202309-11). Written informed consent was obtained from all participants; the study conformed to the Declaration of Helsinki and subsequent amendments.

### Clinical Data Collection and Sample Collection

Baseline demographic characteristics (age, sex, BMI), lifestyle factors (smoking), and comorbidities (hypertension, diabetes duration) were extracted from the hospital’s electronic medical record system. Following overnight fasting, venous blood was drawn on day 2 for routine assays (complete blood count, liver/renal function, lipid panel, FPG, HbA1c), centrifuged ((3,000 × g, 4°C, 15 min), and serum archived at −80°C. Fecal samples were self-collected with sterile disposable kits, transported within 2 hours, and snap-frozen at −80°C.“

### Fecal DNA Extraction

Microbial DNA was isolated from fecal aliquots using the cetyltrimethylammonium bromide (CTAB) method, integrity was checked on 1% agarose gels.

### 16S rRNA Sequencing

The V3–V4 region was PCR-amplified, verified on 2% agarose gels, purified with AMPure XT beads (Beckman Coulter), and quantified by Qubit fluorometry (Invitrogen). Libraries were size-selected on an Agilent 2100 Bioanalyzer, re-quantified with the Kapa Illumina Library Quantification Kit, and sequenced on the NovaSeq 6000 platform (Illumina).

### Microbiome Data Analysis

Fecal DNA underwent paired-end sequencing on Illumina NovaSeq (LC-Bio). Reads were merged via FLASH, trimmed for low-quality bases, filtered with fqtrim (v0.94), and dechimerized with Vsearch (v2.3.4); DADA2 denoising yielded feature tables and representative sequences. Taxonomic annotation was conducted via BLAST alignment to the SILVA 138 database. Post-rarefaction, α/β-diversity indices were computed in QIIME2 and visualized using R (v3.5.2).

### Serum Samples Preparation

Proteins were precipitated from 100 µL serum aliquots with 400 µL ice-cold 50% methanol (−20°C, 30 min), followed by centrifugation (20,000 × g, 15 min). Supernatants were re-centrifuged and analyzed by UPLC-HRMS. QC specimens were prepared by pooling 10 µL of each extract.

### Untargeted Metabolomics Analysis

Metabolites were resolved on an ACQUITY UPLC T3 column (Waters, 40°C) with a 10-min gradient (A: 5 mmol/L ammonium acetate/acetic acid in water; B: acetonitrile; 0.3 mL/min). HRMS detection used a Q-Exactive Plus (Thermo Fisher Scientific, +4.0/–2.8 kV, data-dependent acquisition), with QC samples interspersed every 10 injections for stability monitoring.

### Metabolomic Data Analysis

XCMS-processed mzXML files underwent peak detection, retention time (RT) correction, and isotope annotation to generate an RT–m/z matrix. Metabolite IDs were mapped to KEGG/HMDB. After filtering, imputation, and normalization, partial least squares discriminant analysis (PLS-DA) yielded variable importance in projection (VIP) scores; differential metabolites met *P* < 0.05, |log₂FC| > 0.26, and VIP ≥ 1.

### Diagnostic Model Development and Validation

Candidate biomarkers were selected by the intersection of logistic regression (P < 0.05), LASSO, and random forest analysis using Mean Decrease in Accuracy (RF-MDA). Four diagnostic models—clinical, microbiome, metabolite, and combined—were developed. Model discrimination was quantified by the AUC, sensitivity, and specificity; calibration was assessed using the Hosmer-Lemeshow goodness-of-fit test; and clinical utility was evaluated via decision curve analysis (DCA). Additionally, internal validation was performed using bootstrap resampling (1,000 iterations) and repeated 5-fold cross-validation (10 repetitions) to confirm model robustness.

### Statistical Analysis

Continuous variables were compared via t-tests or Mann-Whitney U tests; categorical variables via χ² tests. Correlation heatmaps illustrating relationships between clinical indicators, microbiota, and metabolites were generated based on Spearman rank correlation analysis. Data analyses used SPSS 26.0 and R 4.2.0; P < 0.05 indicated significance.

## Results

### Baseline Characteristics of the Patients

This study comprised 38 patients with T2DM complicated by ACS (T2DM–ACS group: 27 males, 11 females) and 38 patients with T2DM alone (T2DM group: 30 males, 8 females). Groups were balanced for age and sex (*P* > 0.05). The T2DM-ACS group exhibited lower high-density lipoprotein cholesterol (HDL-C), ejection fraction (EF), absolute lymphocyte count (LYM), and absolute eosinophil count (EOS), with elevated neutrophils (NEUT) and alanine aminotransferase (ALT) (*P* < 0.05, Table 1).

**Table 1.** Baseline Clinical Characteristics of Study Participants.

| Variable | T2DM-ACS (n = 38) | T2DM (n = 38) | P |
| --- | --- | --- | --- |
| Age (years) | 57.39 ± 6.91 | 56.26 ± 6.93 | 0.32 |
| T2DM duration (years) | 9.08 ± 5.44 | 9.05 ± 5.63 | 0.981 |
| BPM (beats/min) | 79.61 ± 12.91 | 76.71 ± 11.22 | 0.486 |
| BMI (kg/m<sup>2</sup>) | 24.78 ± 3.36 | 25.47 ± 2.59 | 0.164 |
| SBP (mmHg) | 135.05 ± 20.77 | 132.45 ± 14.28 | 0.751 |
| DBP (mmHg) | 82.03 ± 11.97 | 84.00 ± 10.29 | 0.443 |
| WBC (10<sup>9</sup>/L) | 6.86 ± 2.03 | 6.11 ± 1.24 | 0.057 |
| LYM (10<sup>9</sup>/L)* | 1.60 ± 0.57 | 2.01 ± 0.43 | **0.003** |
| MONO (10<sup>9</sup>/L) | 0.48 ± 0.20 | 0.47 ± 0.11 | 0.596 |
| NEUT (10<sup>9</sup>/L)* | 4.63 ± 1.66 | 3.45 ± 1.02 | **0.001** |
| EOS (10<sup>9</sup>/L)* | 0.11 ± 0.08 | 0.15 ± 0.12 | **0.037** |
| BASO (10<sup>9</sup>/L) | 0.03 ± 0.02 | 0.03 ± 0.01 | 0.392 |
| ALT (U/L)* | 26.87 ± 15.01 | 19.95 ± 6.45 | **0.011** |
| AST (U/L) | 24.42 ± 9.83 | 22.32 ± 11.99 | 0.176 |
| BUN (mmol/L) | 5.86 ± 1.41 | 6.00 ± 1.50 | 0.675 |
| Cr (μmol/L) | 63.27 ± 22.05 | 63.11 ± 13.71 | 0.232 |
| UA (μmol/L) | 304.36 ± 70.40 | 309.11 ± 62.57 | 0.757 |
| eGFR (ml/min) | 102.79 ± 23.70 | 102.00 ± 12.65 | 0.913 |
| TC (mmol/L) | 1.75 ± 0.69 | 2.01 ± 1.56 | 0.992 |
| TG (mmol/L) | 3.69 ± 1.17 | 4.17 ± 0.98 | 0.055 |
| HDL-C (mmol/L)* | 0.93 ± 0.23 | 1.05 ± 0.21 | **0.016** |
| LDL-C (mmol/L) | 1.99 ± 0.87 | 2.32 ± 0.81 | 0.092 |
| FPG (mmol/L) | 9.10 ± 2.92 | 8.51 ± 3.17 | 0.207 |
| HbA1c (%) | 8.32 ± 1.59 | 8.56 ± 2.08 | 0.909 |
| EF (%)* | 61.37 ± 6.87 | 65.39 ± 3.75 | **0.002** |
| Male (%) | 27/38 | 30/38 | 0.427 |
| Smoking (%) | 15/38 | 16/38 | 0.815 |
| HTN (%) | 19/38 | 16/38 | 0.49 |

\*Indicates statistically significant difference between groups ( $P < 0.05$ ).

### Characteristics of the Gut Microbiota

16S rRNA amplicon profiling of 76 fecal samples revealed 1,754 shared ASVs, 3,200 unique to the T2DM-ACS group, and 2,731 unique to the T2DM group (Fig. S1A). Rarefaction curves plateaued, confirming adequate depth (Fig. S1B). Alpha diversity indices (Chao1, observed species, Goods_coverage, Shannon, Simpson, Pielou_e) did not differ between groups (P > 0.05; Fig. 1A–F, Table S1), indicating comparable richness and evenness. In contrast, beta diversity differed significantly: unweighted UniFrac principal coordinate analysis (PCoA) showed distinct clustering (P < 0.05; PC1 10.61%, PC2 7.56%; Fig. 1G), with non-metric multidimensional scaling (NMDS) corroborating these differences (Fig. 1H).

**Fig. 1.**
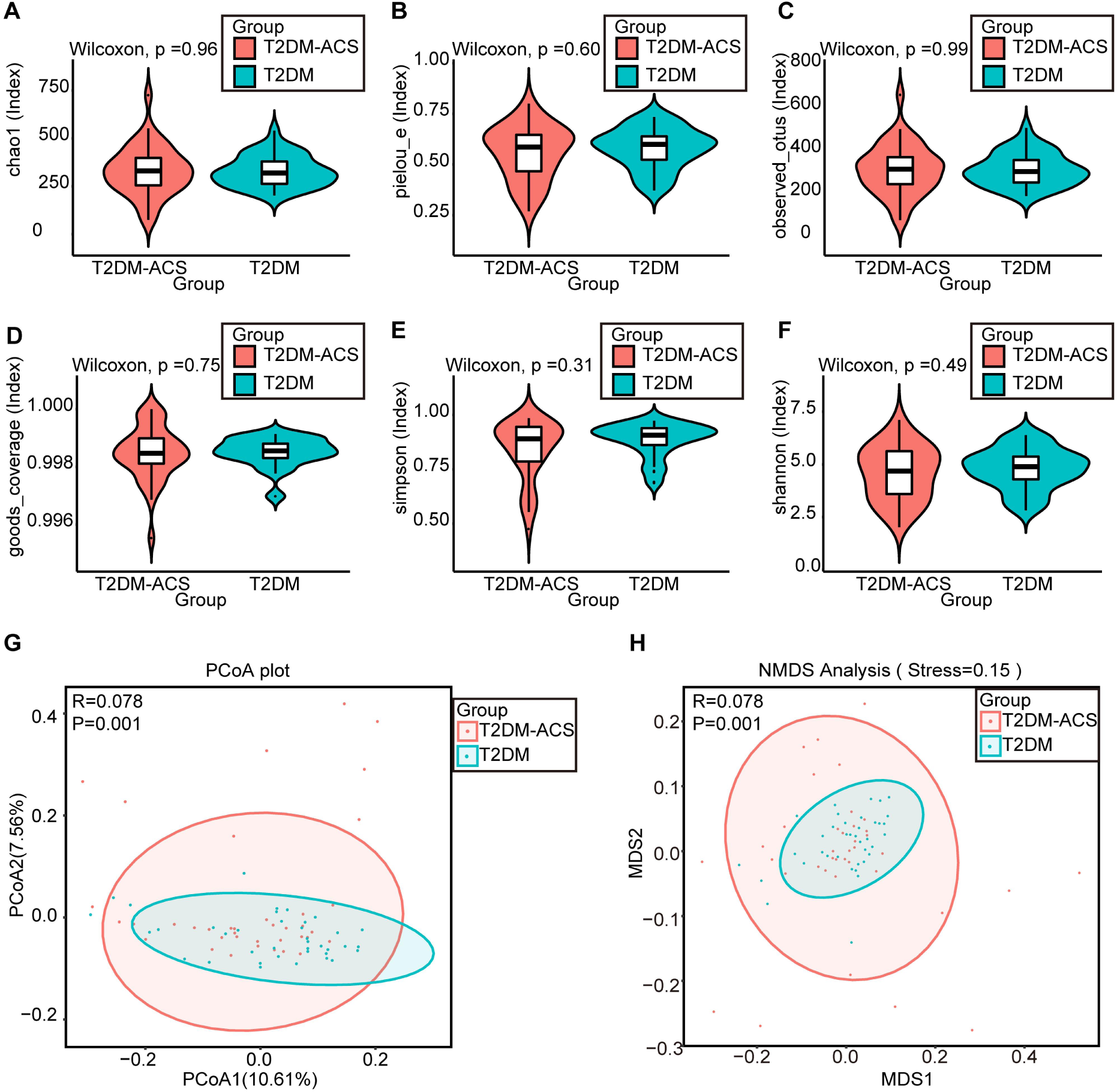
Microbial community diversity comparisons between the T2DM-ACS group (n=38) and T2DM group (n=38). **A–F**: Alpha diversity analyses: **A**, Chao1 index; **B**, Pielou’s evenness index (Pielou_e); **C**, Observed operational taxonomic units (OTUs); **D**, Good’s coverage (Goods_coverage) index; **E**, Simpson index; **F**, Shannon index. Group comparisons were conducted using the Wilcoxon rank-sum test. **G–H**: Beta diversity analyses based on unweighted UniFrac distances: **G**, Principal Coordinates Analysis (PCoA); **H**, Non-metric multidimensional scaling (NMDS) analysis (stress=0.15, indicating satisfactory fit).

*Firmicutes* and *Proteobacteria* predominated in both groups at the phylum level; similarly, Bifidobacterium and Bacteroides were the dominant genera (Fig. 2A–B). Differential abundance analysis revealed 4 discriminatory phyla: *Synergistota* enriched in the T2DM-ACS group, *Cyanobacteria* and *Patescibacteria* depleted (P < 0.05; Fig. S1C). At the genus level, 36 genera exhibited significant differential abundance: 15 were enriched in the T2DM-ACS group (including *Enterococcus*, *Alloprevotella*, and *Butyricimonas*), while 21 were depleted (e.g., *Megasphaera* and *Lachnospira*) (P < 0.05; Fig. 2C, Table S2). These genera were subsequently used for biomarker screening.

**Fig. 2.**
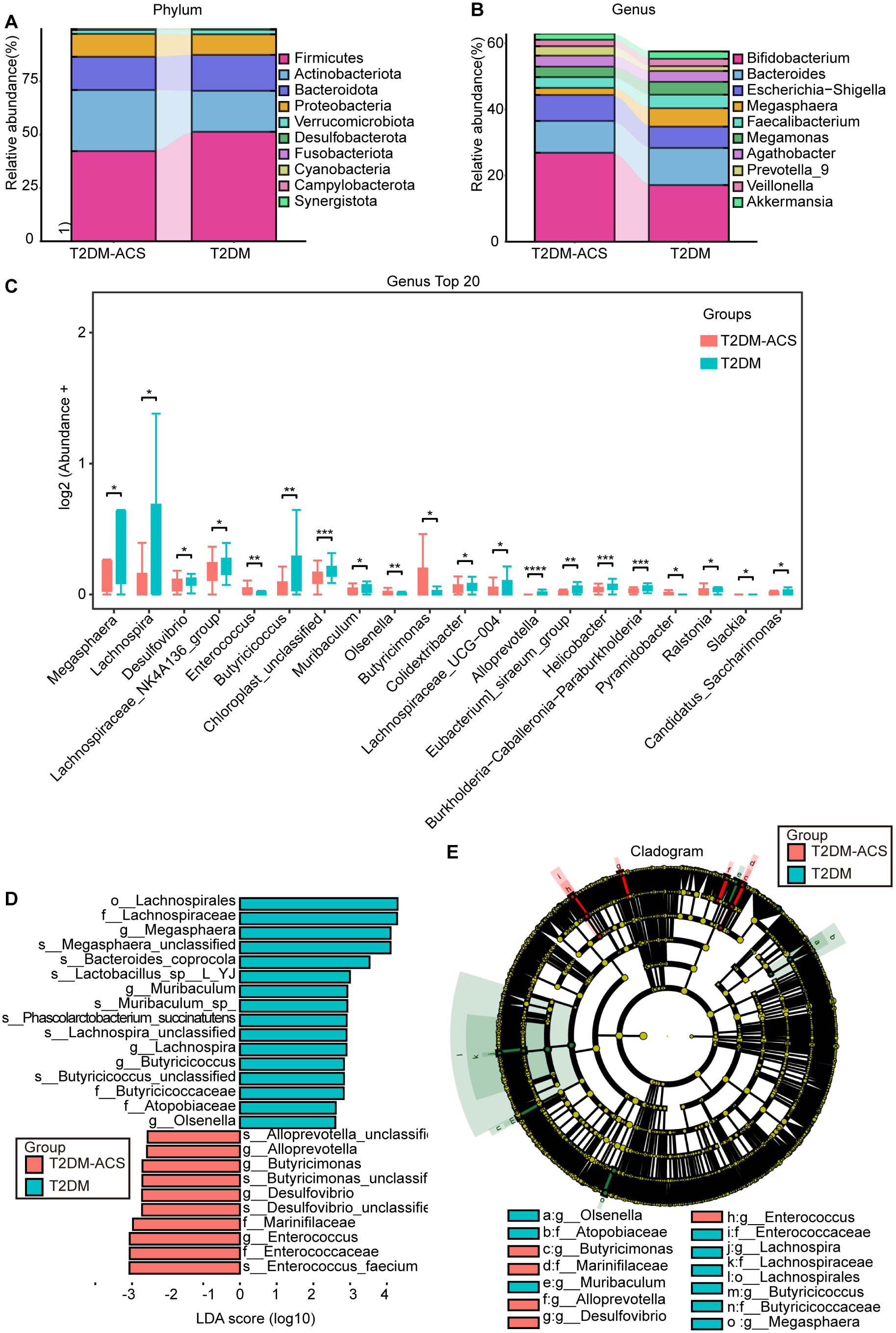
Identification of Differentially Abundant Gut Microbiota Between the T2DM-ACS and T2DM Groups. **A–B**: Stacked bar plots showing the top 10 taxa by relative abundance at the phylum level (**A**) and genus level (**B**); **C**: Intergroup differential analysis of the top 20 genera. \**P* < 0.05, \*\**P* < 0.01, \*\*\**P* < 0.001 (Wilcoxon rank-sum test with false discovery rate [FDR] correction); **D–E**: Linear discriminant analysis effect size (LEfSe) analysis results: **D**: Histogram of linear discriminant analysis (LDA) score distribution (LDA score ≥ 2.5, *P* < 0.05), showing signature species with intergroup differences; **E**: Cladogram, where differently colored nodes indicate species enriched in the corresponding groups, with taxonomic levels labeled (o: Order, f: Family, g: Genus, s: Species).

Linear discriminant analysis effect size (LEfSe) analysis confirmed that *Enterococcus* and *Desulfovibrio* were enriched as biomarkers in the T2DM-ACS group, whereas *Butyricicoccus* and *Lactobacillus* were enriched in the T2DM group (Fig. 2D-E).

### Untargeted Serum Metabolomic Analysis

To probe gut–serum metabolic crosstalk, serum metabolomes were profiled by untargeted LC–MS, yielding 1,056 annotated metabolic features (Table S3).

PLS-DArevealed a clear separation between the T2DM-ACS group and the T2DM group. The permutation test confirmed no overfitting of the data, with R²Y and Q² values of 0.8139 and −0.3879, respectively(Fig. 3A-B).

**Fig. 3.**
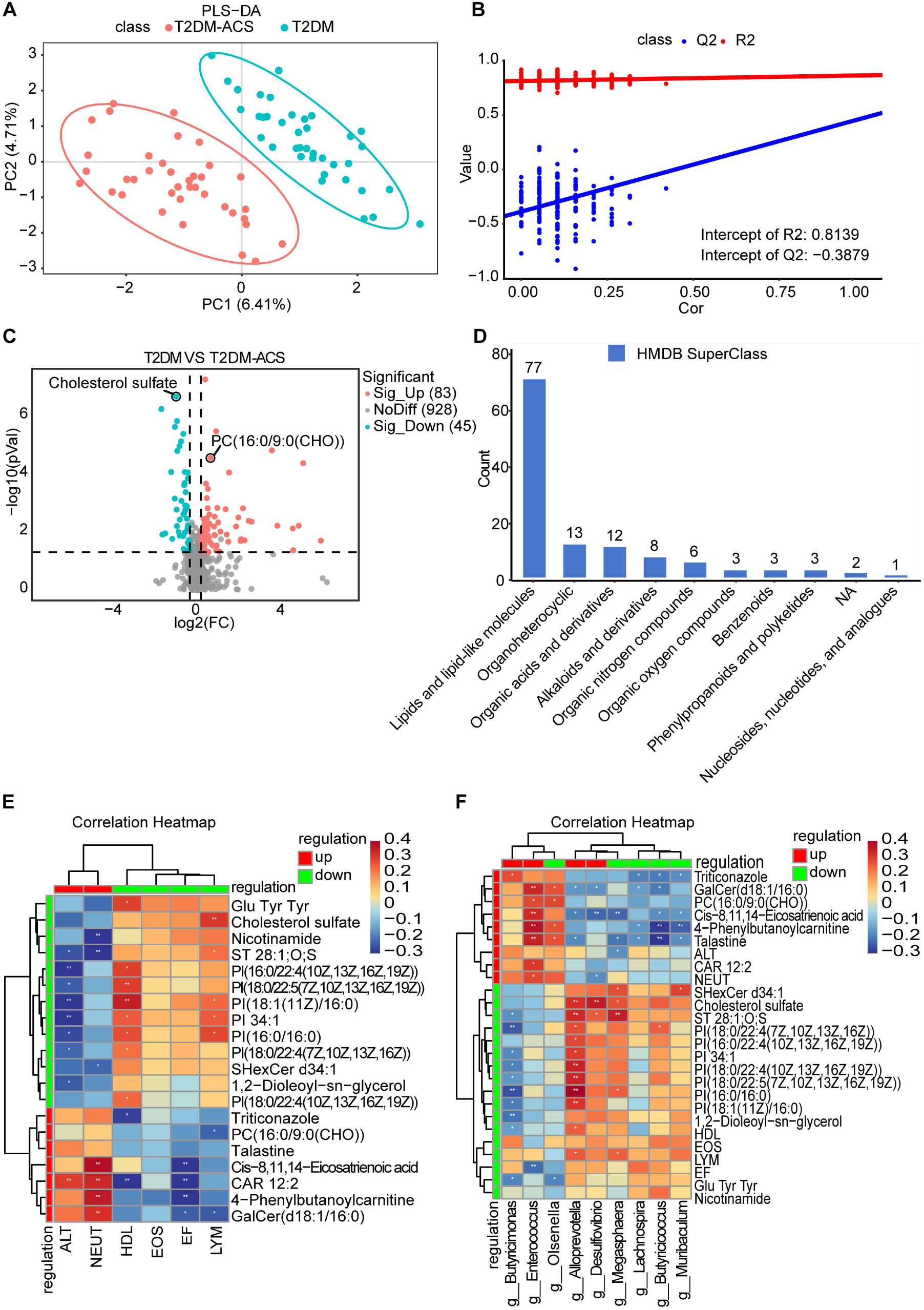
Metabolomic Profiles and Multi-omics Correlation analysis. **A**: PLS-DAscore plot; **B**: PLS-DA permutation test plot; **C**: Volcano plot of differential metabolites. Screening criteria: *P* < 0.05, FC ≥ 1.2 or FC ≤ 1/1.2, and VIP ≥ 1; **D**: Distribution of differential metabolites across Human Metabolome Database (HMDB) superclasses; **E–F**: Correlation heatmaps depicting spearman correlation coefficients among differential metabolites, gut microbes, and clinical indicators. *P < 0.05, **P < 0.01, ***P < 0.001. **Abbreviations**: PLS-DA, Partial least squares discriminant analysis; ALT, alanine aminotransferase; NEUT, absolute neutrophil count; HDL-C, high-density lipoprotein cholesterol; EOS, absolute eosinophil count; EF, ejection fraction; LYM, absolute lymphocyte count; GalCer, galactosylceramide; PC, phosphatidylcholine; PI, phosphatidylinositol; CAR, carnitine.

VIP scores and univariate testing jointly identified 128 differential metabolites between the two groups, with 83 being upregulated and 45 downregulated in the T2DM-ACS group relative to the T2DM group (*P* < 0.05, |log₂FC| > 0.26, and VIP ≥ 1; Fig. 3C).

Superclass classification of metabolites in the HMDB revealed that the differentially abundant metabolites were mainly categorized into lipids and lipid-like molecules, including cholesterol sulfate (downregulated) and PC(16:0/9:0(CHO)) (upregulated). (Fig. 3D).

### Integrated Multiomics Analysis

Spearman’s rank correlation analysis was performed to explore the associations among the screened differential clinical indicators, serum abundant metabolites, and gut microbiota. As shown in Fig. 3E-F, *Enterococcus* was negatively correlated with EF (r = −0.30, *P* < 0.001) and positively correlated with NEUT, PC(16:0/9:0(CHO)), GalCer(d18:1/16:0), CAR 12:2, and cis-8,11,14-eicosatrienoic acid (all *P* < 0.05). *Butyricimonas* was negatively correlated with HDL, ST 28:1;O;S, and multiple phosphatidylinositol-related substances (all *P* < 0.05). In contrast, *Alloprevotella* was positively correlated with these factors (all *P* < 0.05). LYM was positively correlated with cholesterol sulfate but negatively correlated with PC(16:0/9:0 (CHO)) (both *P* < 0.05). These findings suggest that alterations in the abundance of specific genera may be involved in the pathogenesis of ACS in patients with T2DM through the regulation of lipid metabolism disorders, inflammatory activation, insulin resistance, and other pathways. These findings not only reveal the unique metabolic characteristics of T2DM complicated by ACS but also lay a foundation for the subsequent development of biomarkers.

### Screening of Characteristic Biomarkers and Construction of a Disease Prediction Model

Three statistical methods—logistic regression analysis, LASSO regression, and RF—were employed to screen characteristic biomarkers. The detailed screening process is illustrated in Fig. 4. Two clinical indicators (LYM and NEUT), two bacterial genera (*Enterococcus* and *Butyricimonas*), and two serum metabolites (Cholesterol sulfate and PC(16:0/9:0(CHO)) were ultimately included in the model(Fig. 5, Table S4-S6).

**Fig. 4.**
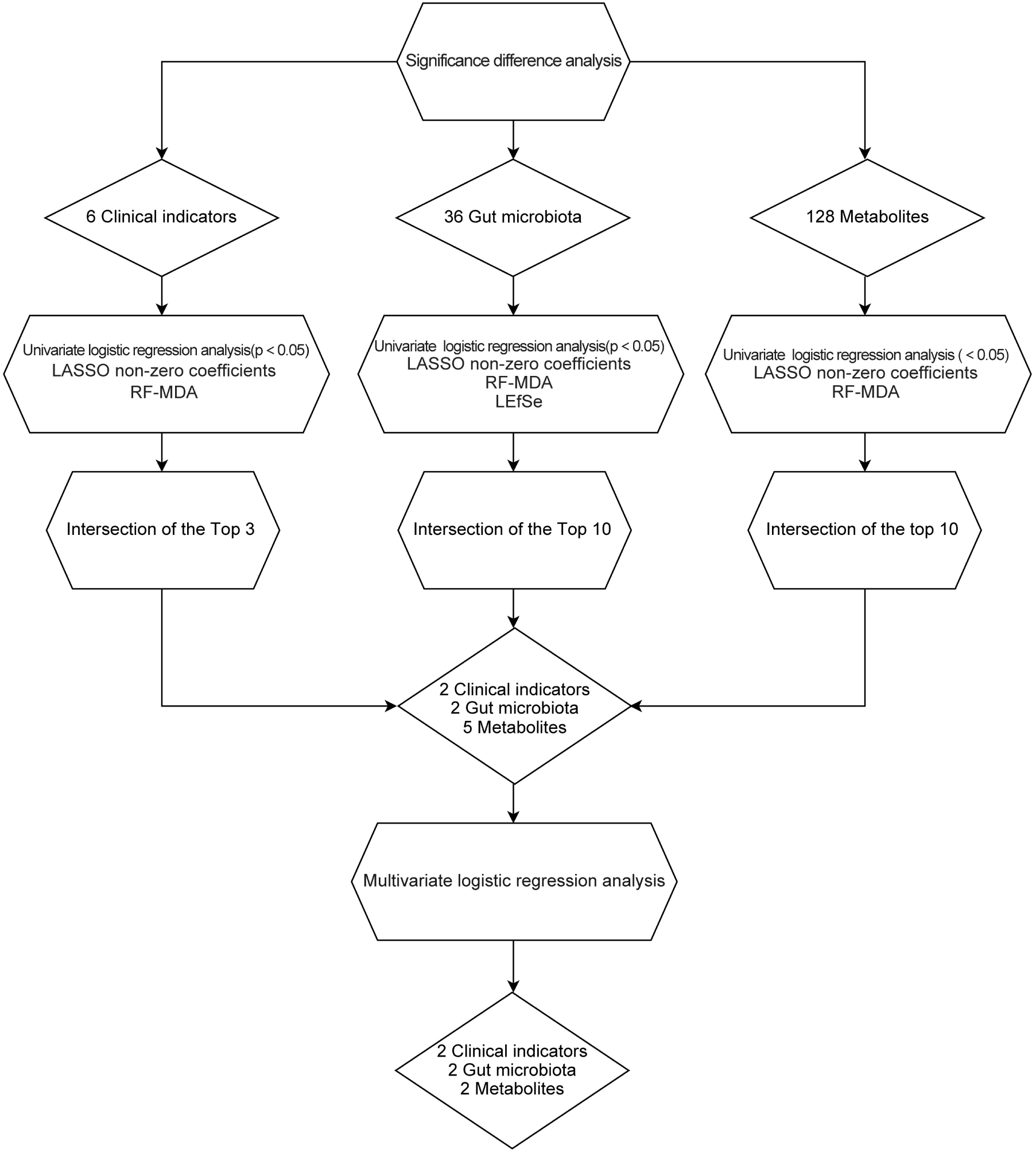
Flowchart of Signature Biomarker Screening for Differentiating T2DM-ACS and T2DM. The analysis pipeline began with significance difference testing of clinical indicators (n=6), gut microbial taxa (n=36), and serum metabolites (n=128) between the two groups. Feature selection was performed using a variety of complementary methods: univariate logistic regression, LASSO, RF-MDA, and LEfSe. The top-ranked features from each method were intersected to identify robust biomarkers. Candidate features (2 clinical indicators, 2 gut microbiota, and 5 metabolites) were then subjected to multivariate logistic regression analysis to construct the final diagnostic model comprising 6 signature biomarkers. **Abbreviations**: LASSO, least absolute shrinkage and selection operator; LEfSe, linear discriminant analysis effect size; RF-MDA, random forest mean decrease in accuracy.

**Fig. 5.**
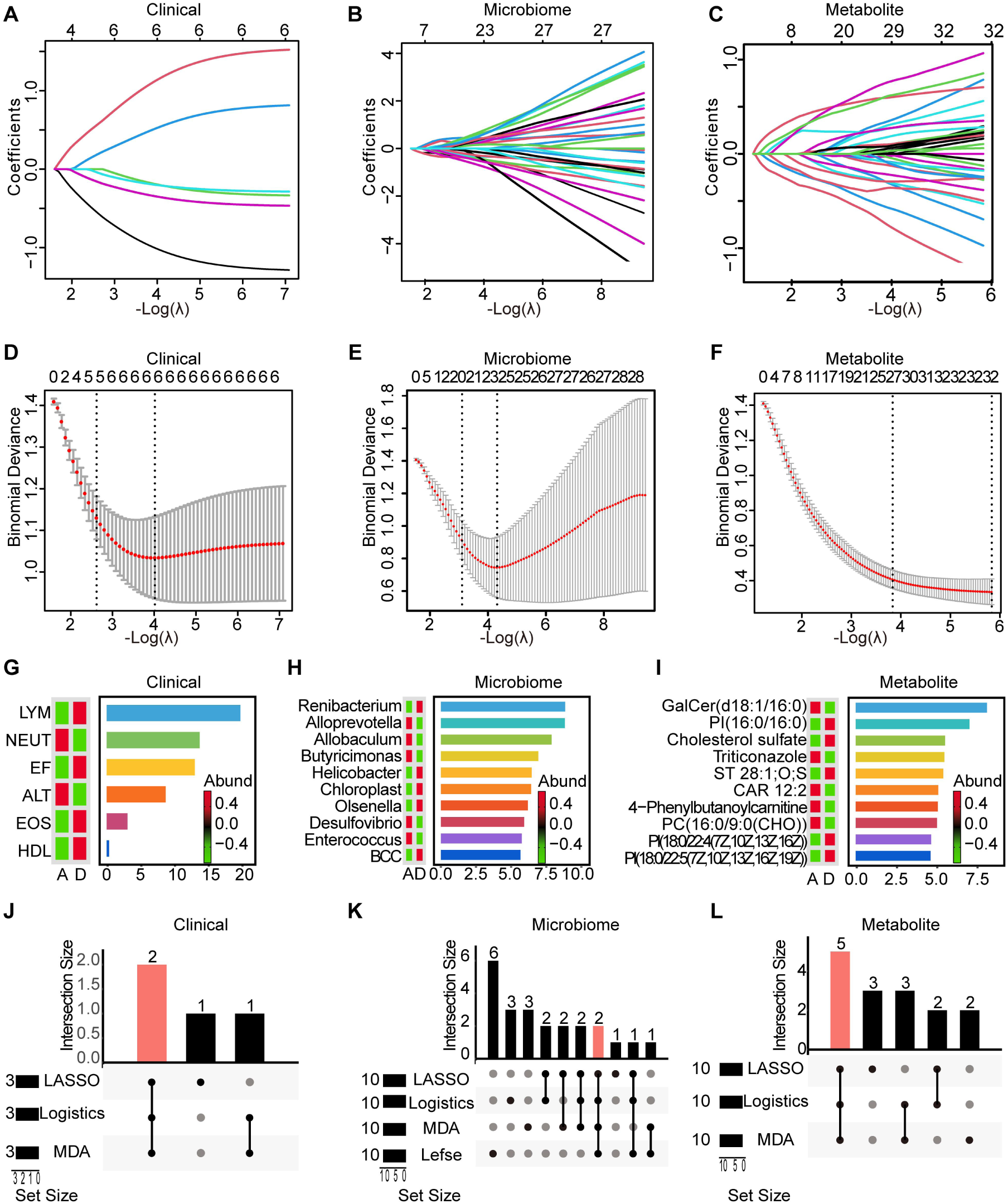
Screening of Characteristic Biomarkers. **A–C**: LASSO coefficient path plot; **D–F**: Cross-validation curve for LASSO regression; **G–I**: MDA values from random forest analysis; **J – L**: Upset plots showing the intersection of features selected by multiple methods. **Abbreviations**: MDA, mean decrease in accuracy; LASSO, least absolute shrinkage and selection operator; LEfSe, linear discriminant analysis effect size; λ, penalty parameter.

To verify their diagnostic value, we constructed four logistic regression models and compared their performance, namely, a clinical model, a microbiome model, a metabolite model, and a combined model (integrating clinical indicators, the gut microbiota, and metabolites). Before incorporating the biomarkers into the combined model, we analyzed and excluded multicollinearity among them. The results revealed that the combined model exhibited the best performance (AUC = 0.983), which was superior to those of the other three individual models (AUC = 0.843, 0.788, and 0.957) (Fig. 6A, Table S7).

**Fig. 6.**
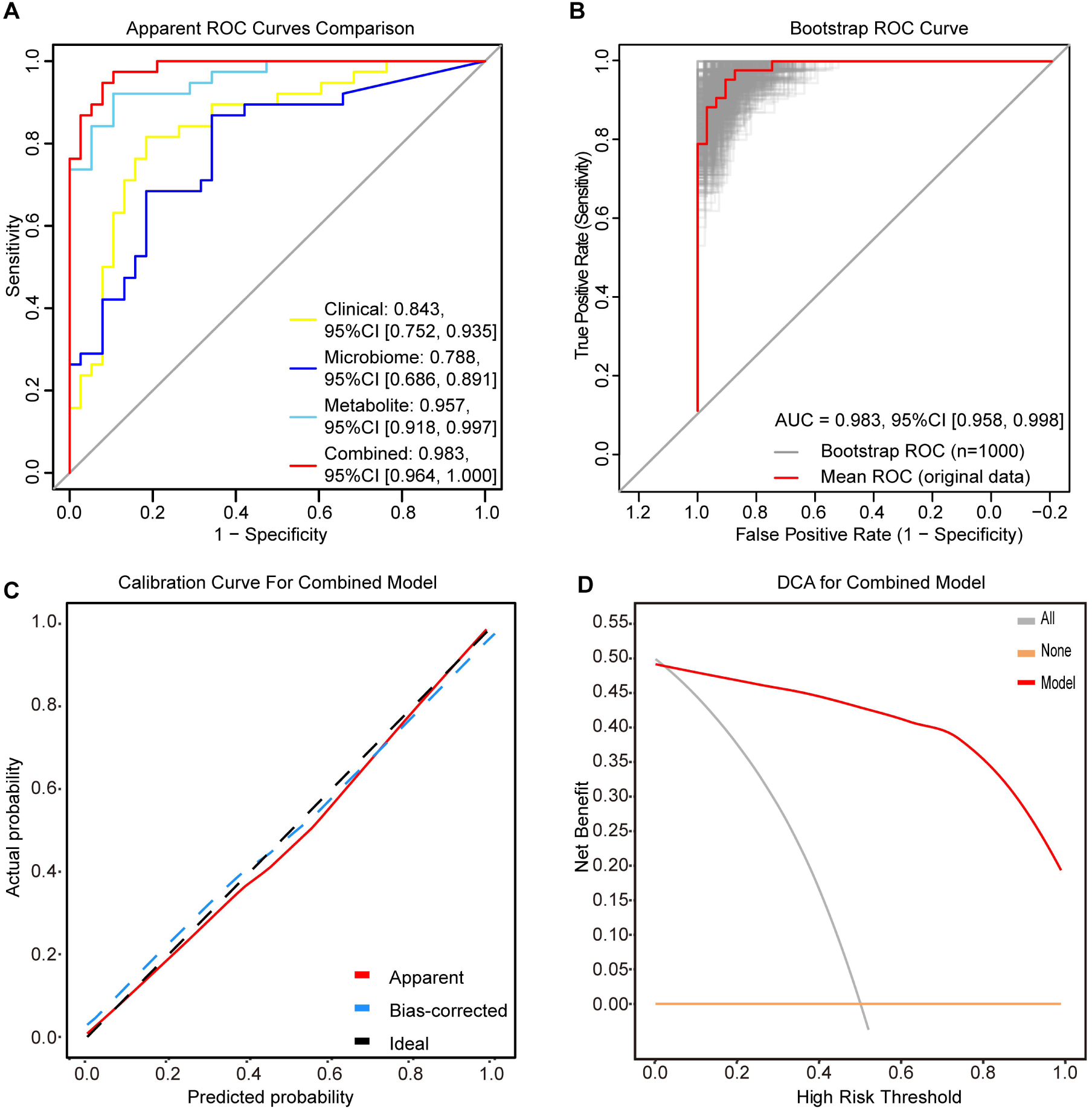
Performance and Validation of the T2DM-ACS Prediction Models. **A**: ROC curves of four prediction models: clinical model, microbiome model, metabolite model, and combined model; **B**:Bootstrap-validated ROC curve of the combined model. Gray lines represent 1000 bootstrap resampling iterations; the red line indicates the mean ROC curve; **C**: Calibration curve of the combined model; **D**: DCA plot of the combined model. **Abbreviations**: AUC, area under the curve; CI, confidence interval; ROC, receiver operating characteristic; DCA, decision curve analysis.

Internal validation of the combined model was subsequently performed. After 1000 bootstrap resamplings, the model’s AUC remained stable at 0.983 (95% CI 0.958–0.998) (Fig. 6B). The results of fivefold cross-validation (repeated 10 times) revealed a slight decrease in diagnostic efficacy (AUC = 0.937). The calibration curve showed minimal deviation between predicted and observed disease probabilities(Fig. 6C). These data demonstrate robust diagnostic performance of the combined model. Decision curve analysis (DCA) further confirmed clinical utility, demonstrating positive net benefit of the combined model across clinically relevant threshold probabilities (Fig. 6D). Notably, the integration of metabolites and the gut microbiota improved the ability to distinguish patients with T2DM with newly diagnosed ACS from those with T2DM alone. Overall, the combination of clinical indicators, gut microbial characteristics, and metabolic profiles is expected to accurately predict disease status in clinical practice.

## Discussion

The increasing global prevalence of T2DM has been paralleled by a disproportionate surge in incident ACS among this population. Consequently, the identification of high-performance biomarkers that permit early risk stratification, timely diagnosis, dynamic surveillance, and precision therapy is imperative. In the present study, we delineated—within a single analytical framework—distinct perturbations in clinical phenotypes, gut microbial architecture, and serum metabolomic signatures that differentiate individuals with uncomplicated T2DM from those with newly diagnosed ACS complicated with T2DM. By integrating these multiomic strata, we uncovered a parsimonious set of biomarkers that herald ACS onset in the context of T2DM. Furthermore, a composite diagnostic algorithm incorporating clinical variables, enteric microbial features, and circulating metabolites achieved commendable discriminatory accuracy for the detection of ACS complicated with T2DM.

Compared with individuals with uncomplicated T2DM, patients with both T2DM and newly diagnosed ACS presented pronounced decreases in the levels of HDL-C, EF, LYM, and EOS, which were concomitant with significant increases in the levels of NEUT and ALT. These perturbations mirror the classical cardiovascular risk profile, underscoring a pathobiology in which dysregulated lipid metabolism and systemic inflammatory activation converge.

Relative neutrophilia and lymphopenia indicate a heightened systemic inflammatory tone that is intimately linked to coronary plaque destabilization and thrombotic propensity in ACS patients^20,21^. Driven by inflammatory cues, neutrophils transmigrate across the activated endothelium, infiltrate nascent plaques, and discharge an arsenal of proinflammatory cytokines that amplify atherogenesis. Moreover, neutrophil extracellular traps (NETs) provoke endothelial injury and paracellular death, thereby leading to plaque rupture. A sterile inflammatory milieu is now recognized as a key driver of acute myocardial infarction and ischemia–reperfusion damage and is characterized by leukocytosis, neutrophilia, and monocytosis. HDL-C, the archetypal atheroprotective lipoprotein, has anti-inflammatory, antioxidant, and anti-diabetogenic capacities. Coronary atherogenesis accelerates when HDL-C is reduced in the context of T2DM^22,23^. Consequently, an elevated neutrophil-to-HDL-C ratio (NHR) has emerged as a readily quantifiable risk indicator for ACS complicated with T2DM^24^.

Gut dysbiosis is intimately involved in the pathogenesis of T2DM and in communication with the cardiovascular system via the gut–heart axis, modulating inflammatory, immune, and metabolic circuits that precipitate CAD^25^. Here, we demonstrate that the fecal microbiota of patients with T2DM who develop ACS undergoes profound structural reconfiguration, a pattern echoing recent reports in CAD, stroke, and chronic heart failure^26–28^. At the genus level, compared with patients with T2DM only, *Enterococcus* and *Butyricimonas* emerged as the most conspicuously enriched taxa in patients with T2DM complicated by ACS.

*Enterococcus* spp. are opportunistic pathogens whose expansion has been reproducibly linked to cardiometabolic complications^29–31^. In a recent murine study, Li et al. reported that enterococcal-derived tyramine activates intestinal epithelial α2A-adrenergic receptors, thereby suppressing Lgr5+ intestinal stem-cell proliferation and compromising barrier integrity^32^. The overgrowth of enterococci may additionally amplify intestinal leakage through lipoteichoic acid and enterotoxin production, facilitating the translocation of lipopolysaccharide and oxidized LDL into the systemic circulation, fuelling endothelial dysfunction and atherogenic inflammation^21,33,34^. Moreover, *Enterococcus faecalis* is a leading causative agent of infective endocarditis, and cohort analyses have revealed a higher incidence of ischemic cardiomyopathy in enterococcal endocarditis than in endocarditis of other etiologies^35^.

*Butyricimonas*, a canonical butyrate producer, generates short-chain fatty acids (SCFAs) that attenuate inflammation, reinforce epithelial barrier function, and fine-tune glucolipid metabolism^36,37^. In ApoE−/− atherosclerotic mice, Tian et al. demonstrated that exogenous butyrate restored endothelial function via PPARδ/miR-181b signaling and attenuated plaque formation^38^. Paradoxically, we observed a significant expansion of *Butyricimonas* in patients with T2DM complicated by ACS, a finding concordant with recent observations in acute myocardial infarction^20^, CAD^31,39^ and a validated primate ischemia–reperfusion model encompassing 77 patients with ST-elevation myocardial infarction and eight rhesus monkeys^40^. We therefore propose that the post-ischemic enrichment of *Butyricimonas* represents a compensatory reparatory response aimed at re-establishing microbial homeostasis and mitigating cardiac injury after ACS. Our dataset provides robust translational evidence for the contextual role of *Butyricimonas* in the gut–heart axis and underscores the necessity of dissecting its functional attributes under hyperglycemic stress.

Concomitant with the observed gut microbial dysbiosis, we documented a characteristic serum metabolomic shift in patients with T2DM complicated by ACS. Compared with T2DM controls, the ACS group exhibited significant upregulation of PC(16:0/9:0(CHO)) and arachidonic acid and downregulation of cholesterol sulfate (CS); collectively, these discriminating signals belong to the lipid and lipid-like molecular superclass.

Phosphatidylcholine (PC), the quantitatively dominant phospholipid in mammalian plasma, constitutes the principal precursor pool for AA and lysophosphatidylcholine (LysoPC). These bioactive lipids function as potent inflammatory mediators that orchestrate insulin resistance and drive the initiation, progression, and ultimately rupture of atherosclerotic plaques^41,42^. In a cohort of 101 individuals, Ménégaut et al. demonstrated significant enrichment of both AA and LysoPC in plasma and carotid plaque specimens from T2DM patients with established atheroma^43^, corroborating earlier epidemiological data that identified circulating PC as an independent predictor of CAD^44^. Mechanistic support is provided by Cole et al., who showed that genetic or pharmacologic impairment of hepatic PC biosynthesis attenuates atherosclerotic burden and improves cardiac performance in hyperlipidemic mice^45^. In addition to its canonical role as a membrane constituent, dietary PC can be channeled by the intestinal microbiota into trimethylamine, which is subsequently oxidized to trimethylamine-N-oxide (TMAO). TMAO amplifies macrophage scavenger receptor expression, impairs reverse cholesterol transport, and incites endothelial inflammation, thereby accelerating atherogenesis^46–48^.

Multiple studies have shown that in patients with diabetes, plasma TMAO can disrupt glucose homeostasis, leading to insulin resistance, and represents a major risk factor for cardiovascular complications^48–52^. Currently, TMAO has been confirmed as an important diagnostic biomarker for ACS^53^.

Cholesterol sulfate, the predominant circulating sulfated steroid, modulates glucolipid homeostasis and engages in bidirectional crosstalk with the gut microbiota^54^. Exogenous CS administration mitigates hyperglycemia in diabetic rodents^55^, attenuates leukotriene biosynthesis, and exerts systemic anti-inflammatory effects^56^. Recent in vitro and in vivo work by Nam et al. revealed that CS lowers intracellular cholesterol by antagonizing its synthesis and uptake pathways while simultaneously diminishing LDL-cholesterol influx^57^. The selective depletion of CS observed in our T2DM–ACS cohort may therefore eliminate a physiological brake on inflammation and lipid accumulation, synergizing with hyperglycemia to precipitate acute coronary events.

Current algorithms for the early triage of ACS in patients with T2DM remain suboptimal, prompting an urgent demand for high-performance biomarkers that can accurately flag incident events and thereby attenuate the population-level cardiovascular burden. Although technological advances have expanded the biomarker repertoire to include cardiomyocyte-restricted proteins (e.g., heart-type fatty acid-binding protein), leukocyte-derived enzymes (e.g., myeloperoxidase), neurohypophyseal hormones (e.g., copeptin) and circulating microRNAs, their incremental diagnostic value in the hyperacute phase is still limited^58–60^. Recent studies have indicated that the algorithmic fusion of heterogeneous biomolecular layers with machine-learning architectures can markedly **i**ncrease discriminative accuracy^61^.

Capitalizing on this conceptual advance, we deployed a nested pipeline comprising LASSO feature selection, RF ensemble learning and logistic regression to identify a parsimonious biosignature capable of discriminating between ACS complicated with T2DM. The input features spanned three orthogonal data modalities: (i) granular clinical phenotypes, (ii) genus-level gut microbial profiles derived from 16S rRNA gene sequencing and (iii) targeted serum metabolome coverage. Individually, each omic stratum manifested respectable discriminatory capacity (clinical-only AUC = 0.983; microbiome-only AUC = 0.788; and metabolome-only AUC = 0.957). However, integrated modeling that allowed cross-omic feature interaction increased the AUC to 0.983, improved precision–recall metrics commensurately, and had calibration curves that demonstrated excellent agreement between the predicted and observed event rates. This systems-level approach not only furnishes a robust quantitative scaffold for early risk stratification but also highlights the therapeutic tractability of microbial and metabolic nodes that drive coronary atherothrombosis^62^. Importantly, our cohort was restricted to newly diagnosed ACS patients, thereby minimizing pharmacological and disease-progression confounders that could otherwise distort the microbiota and metabolite read-outs.

### Limitations

Several caveats merit consideration. First, the modest sample size and single-center recruitment restricted to northwestern China reduce the generalizability of the study results; large-scale, multicenter cohorts are therefore warranted for external validation. Second, although robust associations between specific microbial taxa, metabolites, and ACS were identified, the directionality and causality of these relationships remain unclear. Experimental paradigms such as fecal microbiota transplantation or targeted metabolite supplementation in preclinical models and clinical settings are needed to establish causal inference and to refine patient-selection strategies for T2DM–ACS. Third, the cross-sectional design precludes an appraisal of temporal dynamics; longitudinal follow-up or randomized controlled trials are indispensable for corroborating our observations and delineating causal pathways. Fourth, potential confounders—including dietary habits, physical activity, and pharmacotherapy for diabetes or ACS—were not incorporated into the analytical framework and should be addressed in future investigations. Finally, the reliance on 16S rRNA gene profiling provides only genus-level resolution and limited functional annotation; shotgun metagenomic sequencing will be necessary to obtain a comprehensive catalog of microbial genes and pathways implicated in disease pathophysiology.

## Conclusion

In conclusion, our study revealed a distinct signature of gut dysbiosis and serum metabolomic chaos in patients with T2DM complicated with newly diagnosed ACS. Clinical phenotypes, the fecal microbiome, and circulating metabolites do not operate as isolated entities; instead, they constitute a densely interconnected network in which inflammatory amplification loops synergistically accelerate atherothrombotic vulnerability in the diabetic vasculature. Integrative interrogation of these three data dimensions therefore reconstructs a more complete pathophysiological landscape and illuminates the mechanisms through which T2DM progresses to ACS. The translational utility of this multiomics panel is underscored by a diagnostic model that achieves robust discriminatory performance, implying that a composite index incorporating clinical indices, microbial taxa, and lipid-derived metabolites could serve as an early warning system for ACS onset in patients with T2DM. These findings provide novel insights and potential translational strategies for the clinical surveillance and personalized treatment of patients with T2DM complicated by ACS. Future efforts should prioritize large-scale, multiethnic validation cohorts and randomized trials that formally test whether microbiota-directed interventions can reduce cardiovascular complications in this high-risk population.

## Nonstandard Abbreviations and Acronyms

T2DM: Type 2 diabetes mellitus
ACS: acute coronary syndrome
CAD: coronary artery disease
QC: quality control
ROC: receiver operating characteristic
AUC: area under the curve
TMAO: trimethylamine-N-oxide
LEfSe: Linear discriminant analysis effect size
LC-MS: liquid chromatography-mass spectrometry
LASSO: least absolute shrinkage and selection operator
DCA: decision curve analysis
RF-MDA: random forest analysis using Mean Decrease in Accuracy

## Acknowledgements

The authors thank the staff and participants of the Project in Tangdu Hospital for their important contributions.

## Sources of Funding

The work is supported by the research grants from National Natural Science Foundation of China (No. 82270366) and Shaanxi Province Key Research and Development Project (No. 2024GH-ZDXM-40, No. 2025SF-YBXM-414).

## Disclosures

None.

## Conflicts of Interest

No potential conflicts of interest relevant to this article were reported.

## Footnotes

Supplemental Material is available at Supplemental Material. For Sources of Funding and Disclosures, see page 19.

## Supplemental Material

Tables S1–S7

Figures S1

## Notes

### Competing Interest Statement

The authors have declared no competing interest.

